# The absence of superinfection exclusion of Borna disease virus 2 maintains genomic polymorphisms in persistently infected cells

**DOI:** 10.1101/2024.04.15.589485

**Authors:** Takehiro Kanda, Pauline Dianne Santos, Dirk Höper, Martin Beer, Dennis Rubbenstroth, Keizo Tomonaga

**Affiliations:** Laboratory of RNA viruses, Department of Virus Research, Institute for Life and Medical Science, Kyoto University; Department of Molecular Virology, Graduate School of Medicine, Kyoto University; Institute of Diagnostic Virology, Friedrich-Loeffler-Institute; Department of Mammalian Regulatory Network, Graduate School of Biostudies, Kyoto University

## Abstract

Viruses belonging to the genus *Orthobornavirus* within the family *Bornaviridae* are known to infect various vertebrate species, including mammals and birds. Within the genus, the species *Orthobornavirus bornaense* includes two mammalian viruses: Borna disease virus 1 (BoDV-1), the prototype of the family, and its closely related virus, BoDV-2. BoDV-1 was identified as the causative agent of Borna disease (BD) in horses, sheep, humans, and other mammals. BoDV-2 was first detected in a pony in eastern Austria in 1999 that exhibited severe and incurable neurological symptoms. Although BoDV-2 shares approximately 80% nucleotide identity with BoDV-1, its virological properties, including host range, replication ability, and pathogenicity, remain unclear. In this study, we aimed to investigate the virological properties of BoDV-2 by re-evaluating its whole-genome sequence using RNA sequencing. Compared to the published reference sequence, we identified two nonsynonymous nucleotide substitutions in the large (L) gene. One of these substitutions was found to be critical for the restoration of polymerase activity, enabling the successful recovery of recombinant BoDV-2 (rBoDV-2) through reverse genetics. We also identified two nonsynonymous single-nucleotide polymorphisms (SNPs) in the L gene and one in the phosphoprotein (P) gene. Substitution of these SNPs significantly enhanced the growth ability of rBoDV-2. In addition, our studies showed that BoDV-2 does not induce superinfection exclusion in cells, allowing persistence of low-fitness genome variants for an extended period of time. These findings help to characterize the virological properties of BoDV-2 and shed light on how bornaviruses maintain genetic diversity in infected cells.

**Importance:** Mammalian bornaviruses are a general term for viruses belonging to the genus *Orthobornavirus* that infect mammalian species, e.g., Borna disease virus 1 and 2 (BoDV-1 and -2) and variegated squirrel bornavirus 1 (VSBV-1). Although BoDV-1 and VSBV-1 are reported to be associated with fatal encephalitis in humans, the infectivity and pathogenicity of BoDV-2 in humans remain unclear. To determine the virological properties of BoDV-2, we developed a reverse genetics system for BoDV-2. By using recombinant BoDV-2s, we identified several nucleotides that affect the growth ability of BoDV-2 and revealed the molecular mechanisms through which BoDV-2 maintains genetic heterogeneity in persistently infected cells. This reverse genetics system will accelerate the biological studies of BoDV-2 and contribute to the development of countermeasures against mammalian bornaviruses.

## Introduction

Viruses of the genus *Orthobornavirus* within the family *Bornaviridae* have been documented to infect a wide range of vertebrate species, including mammals and avians (1–3). Currently, the genus comprises 25 viruses belonging to 9 different viral species (4). Among these, *Orthobornavirus bornaense* consists of two mammalian viruses: Borna disease virus 1 (BoDV-1), the prototype of the family *Bornaviridae*, and BoDV-2, a genetically closely related virus to BoDV-1. BoDV-1 has been identified as the etiological agent of Borna disease (BD), a nonpurulent encephalomyelitis affecting horses, sheep, and other mammalian species (1, 2). Conversely, BoDV-2 was only isolated from a single case of a pony showing severe and incurable neurological symptoms in eastern Austria in 1999, where BoDV-1 was not endemic (5). Although the genome sequence of BoDV-2 isolate No/98 shares approximately 80% nucleotide identity with that of BoDV-1 strains (6), its virological properties, including host range, replication ability, and pathogenicity, have not been fully elucidated.

The pathogenicity of BoDV-1 in humans was controversial until two case studies were reported in 2018; these studies demonstrated the presence of the BoDV-1 antigen and RNA in a healthy young man and in two organ transplant recipients who died of encephalitis (7, 8). Subsequently, several studies have reported that infection with BoDV-1 was associated with more than forty fatal cases of encephalitis in humans (9–15). Additionally, variegated squirrel bornavirus 1 (VSBV-1; species *Orthobornavirus sciuri*) was isolated from squirrel breeders who died of encephalitis (16). Therefore, both BoDV-1 and VSBV-1 are zoonotic pathogens that cause fatal encephalitis in humans (17). However, the potential of BoDV-2 to infect and cause disease in humans remains unclear. Considering that BoDV-2 isolate No/98 was isolated from a pony that displayed fatal neurological disorders and that BoDV-2 is genetically more closely related to BoDV-1 than VSBV-1, it is likely that BoDV-2 may also possess zoonotic potential.

The reverse genetics system is a powerful tool for elucidating viral characteristics such as replication and pathogenesis. The reverse genetics of BoDV-1 was initially described by Schneider et al. in 2005; a plasmid expressing the full-length BoDV-1 antigenome and three helper plasmids expressing the nucleoprotein (N), phosphoprotein (P), and large protein (L) encoding the RNA-dependent RNA polymerase were transfected into cells expressing T7 polymerase (18). This system has been further improved by employing highly transfectable HEK293T cells for plasmid transfection and incorporating two helper plasmids expressing matrix protein (M) and glycoprotein (G) (19–21). Given that the genomic components of BoDV-2 closely resemble those of BoDV-1, it is predicted that recombinant BoDV-2 (rBoDV-2) can be easily rescued by employing the same strategy as used for BoDV-1. However, although the complete genome sequence of BoDV-2 has been deposited in NCBI GenBank, previous studies have raised concerns about potential errors in the L gene, indicating the need to reassess the genome sequence of BoDV-2 (6).

In this study, we aimed to investigate the virological properties of BoDV-2 using recombinant viral techniques and therefore redetermine the genome sequence of BoDV-2 isolate No/98, which has been maintained in persistently infected cells. We found two nonsynonymous nucleotide substitutions in the L gene compared to the published sequence. One of these substitutions was crucial for restoring the polymerase activity of L, allowing successful recovery of rBoDV-2 through reverse genetics. Additionally, we identified two nonsynonymous single-nucleotide polymorphisms (SNPs) in the L gene and one in the P gene. Substituting these SNPs significantly enhanced the growth ability of rBoDV-2. Furthermore, our study demonstrated that BoDV-2 does not induce superinfection exclusion in cells, leading to long-term maintenance of low-fitness genome variants in persistently infected cells.

## Results

### Reassessment of the complete genome sequence of BoDV-2

The complete genome sequence of BoDV-2 isolate No/98 is deposited in NCBI GenBank; however, it is suspected that this sequence might contain several errors in the L gene because three blocks of amino acid differences are unnaturally accumulated at the C-terminal domain (CTD) of L compared to those of representative strains of BoDV-1, such as He/80 and V (Fig. 1B) (6). Therefore, we reassessed the whole-genome sequence of BoDV-2 isolate No/98 via RNA sequencing (RNA-seq) using total RNA extracted from persistently infected Vero cells available at the Friedrich-Loeffler-Institut. Compared to the published sequence (accession no. AJ311524), the reassessed sequence contains one and five nucleotide differences in the P and L genes, respectively (Fig. 1A). Two of them in the L gene, uracil at positions 2762 and 3334, which are located in the polyribonucleotidyltransferase (PRNTase) domain (Fig. 1B), were completely substituted with guanine, resulting in amino acid changes of leucine at position 921 with arginine and cysteine at position 1112 with glycine (Fig. 1A). In addition, three SNPs were detected in the L gene, two of which, positions 3743 and 4250, induce nonsynonymous amino acid changes in the CTD of L, though they do not overlap with the blocks of amino acid differences described in a previous paper (Fig. 1A and B) (6). A nonsynonymous SNP was also detected at position 456 of the P gene (Fig. 1A). Furthermore, we performed rapid amplification of cDNA ends (RACE) analysis to determine the exact 3’ and 5’ terminal sequences, which could not be reconstituted by RNA-seq analysis. Similar to BoDV-1 (18, 22), BoDV-2 possesses four nucleotides overhung at the 3’ end of both the genome and antigenome without a complementary sequence at the 5’ end of the opposite strand (Fig. 1C). Compared to the published sequence, the reassessed sequence lacks uracil at the 5’ end of the genome (Fig. 1D).

**Fig. 1.**
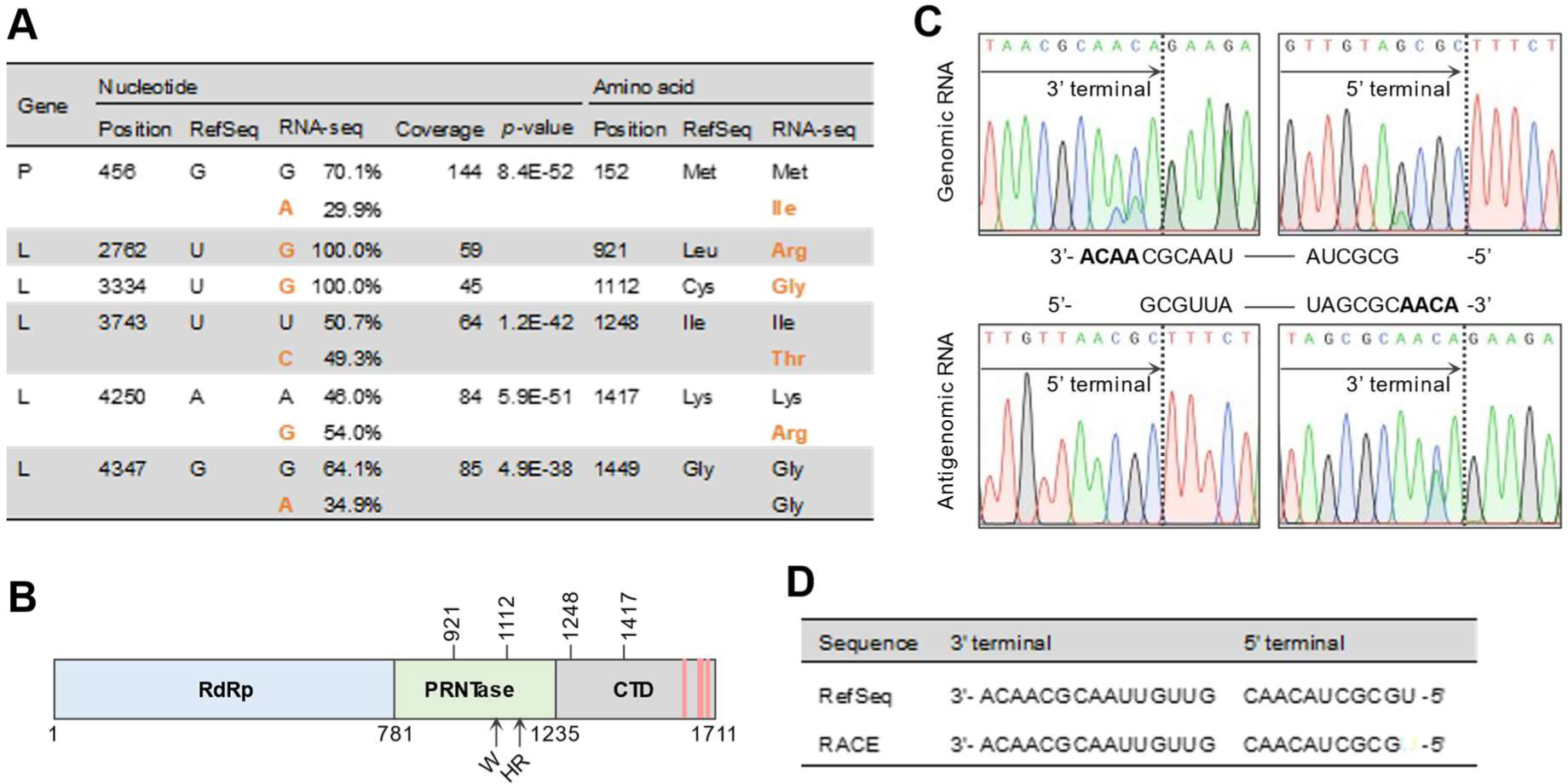
Reassessment of the complete genome sequence of BoDV-2. (A) RNA-seq analysis of total RNA extracted from BoDV-2 isolate No/98-infected Vero cells. Compared to the reference sequence (RefSeq) of BoDV-2 (accession no. NC_030692), substituted nucleotides in the reassessed sequence are represented in bold orange, and their percentages of the total are indicated. Concomitant amino acid changes are also represented in bold orange. (B) Schematic representation of BoDV-2 L. The putative RNA-dependent RNA polymerase (RdRp) domain, polyribonucleotidyltransferase (PRNTase) domain, and C-terminal domain (CTD) are colored blue, green, and gray, respectively. Blocks of amino acid differences between BoDV-1 L and BoDV-2 L in the CTD, which were described in the previous study (6), are represented with pale red bars. The positions at which amino acid changes occur between the reassessed and reference sequences are indicated above, and putative motif C (W) and motif D (HR) in the PRNTase domain are indicated below. (C) RACE analysis of the 5’ and 3’ terminal sequences of both the genome and antigenome of BoDV-2. The 5’ and 3’ ends of total RNA extracted from BoDV-2-infected Vero cells were ligated with each oligoRNA. BoDV-2-specific 5’ and 3’ terminal sequences were amplified via RT-PCR and analyzed via direct sequencing. The border between the viral terminal sequence and ligated oligoRNA is indicated by the dotted line. The analyzed terminal sequences are indicated schematically. (D) Comparison of the reassessed genomic terminal sequences of BoDV-2 with the reference sequence.

### The polymerase activity of BoDV-2 L is restored by substituting a nucleotide

In the reassessed sequence, nucleotides at positions 2762 and 3334 of the L gene are completely substituted compared to the reference sequence (Fig. 1A). To determine whether these differences affect BoDV-2 L gene expression, we constructed L cDNA expression plasmids possessing each nucleotide substitution independently (BoDV-2 L_R_ or L_G_) or in combination (BoDV-2 L_RG_) and performed western blotting (Fig.2A). In a previous study, we could not detect expression of FLAG-tagged BoDV-2 L with the reference sequence, though that of FLAG-tagged BoDV-1 L was clearly demonstrated, suggesting the low expression potential of the FLAG-BoDV2 L construct (23). However, in the present study, we could detect the expression of BoDV-2 L with the reference sequence when a linker sequence was inserted between FLAG and the L gene to enhance the accessibility of the antibody to the tag (Fig. 2B). In addition, we confirmed expressions of the reconstructed BoDV-2 L variants in transfected cells (Fig. 2B).

**Fig. 2.**
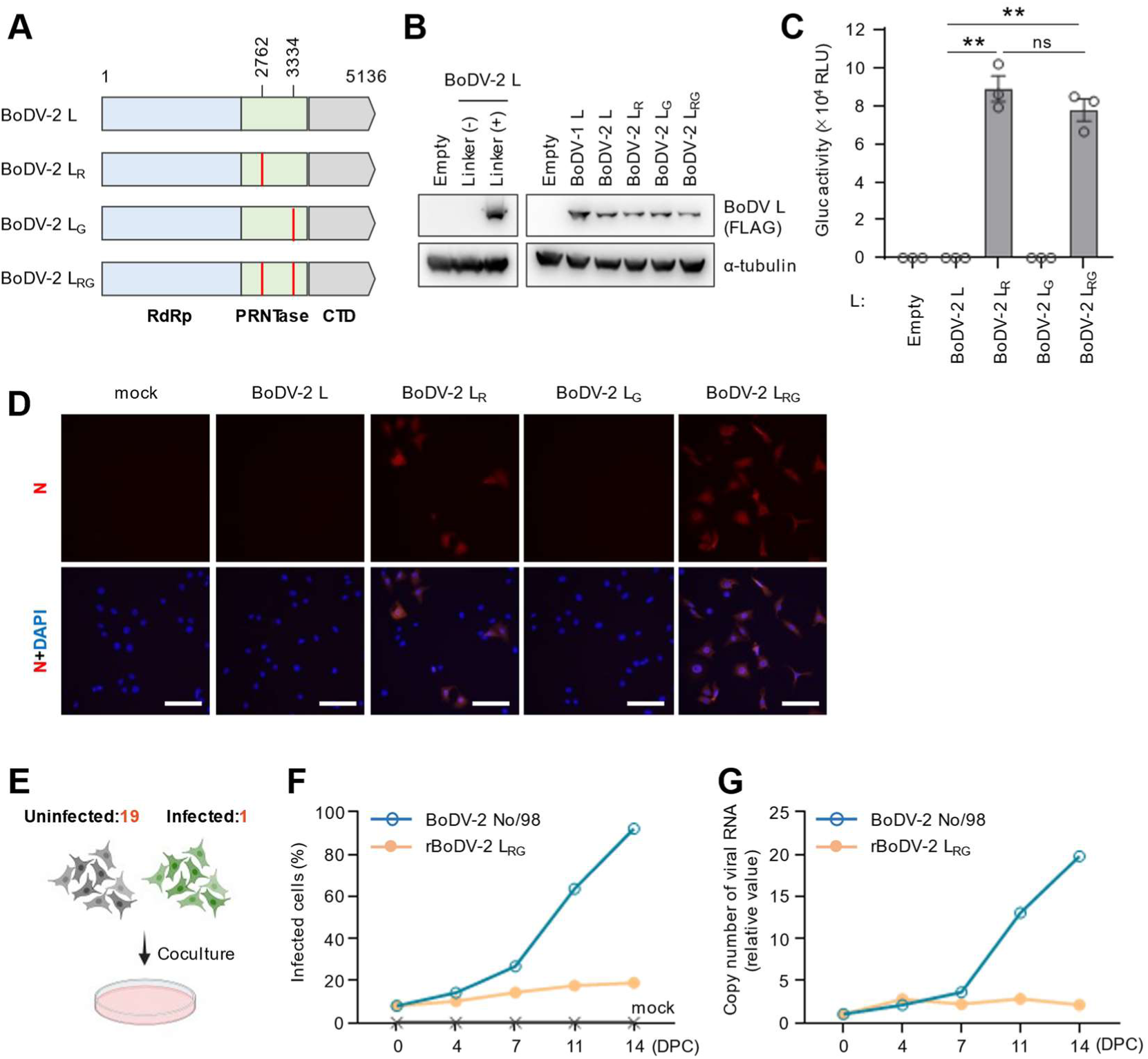
The polymerase activity of BoDV-2 L is restored by substituting a nucleotide. (A) Schematic representation of the BoDV-2 L gene variants inserted into the expression plasmid and the full-length BoDV-2 cDNA plasmid. The positions of the substituted nucleotides compared to the reference sequence are indicated by red bars. The putative RdRp domain, PRNTase domain, and CTD are colored blue, green, and gray, respectively. (B) Detection of BoDV-2 L variants by western blotting analysis. Whole-cell lysates were prepared from 293T cells transfected with the indicated plasmid. The primary antibodies used for detection are indicated to the right of the panels. Alpha-tubulin was used as the loading control. (C) BoDV-2 minireplicon assay using the indicated BoDV-2 L variants and BoDV-2 N and P as helper plasmids. Data are presented as the means ± standard errors of the means (± SEM) of three biologically independent experiments. One-way analysis of variance (ANOVA) and Tukey’s multiple comparisons test were performed for statistical analysis. ns, not significant; **, *p* < 0.01; Gluc, *Gaussia* luciferase; RLU, relative light unit. (D) Detection of rBoDV-2 by immunofluorescence assay (IFA). Reverse genetics was performed by transfecting a BoDV-2 cDNA plasmid possessing the indicated substitution in the L gene and five helper plasmids into HEK293T cells. At 3 days post transfection, transfected HEK293T cells were cocultured with uninfected Vero cells. After 14 days of coculture and complete removal of transfected HEK293T cells, Vero cells were subjected to IFA using an antibody against BoDV N. Nuclei were counterstained with DAPI. Bar, 500 μm. (E to G) Growth kinetics assay of rBoDV-2. (E) Schematic representation of the growth kinetics assay. Vero cells infected with BoDV-2 isolate No/98 or with rBoDV-2 L_RG_ were cocultured with uninfected Vero cells at a ratio of 1 to 19 and passaged every 3 or 4 days. (F) The percentage of infected cells was determined by IFA using an antibody against BoDV N. DPC, days post coculture. (G) The copy number of viral RNA was measured via RT-qPCR using total RNA extracted from each cells. DPC, days post coculture.

To determine the polymerase activities of the reconstructed BoDV-2 L variants, we performed a BoDV-2 minireplicon assay. As shown in Fig. 2C, reporter activity was not detected for BoDV-2 L with the reference sequence, indicating that it lacks any polymerase activity. Interestingly, BoDV-2 L possessing a nucleotide substitution at position 2762, namely, BoDV-2 L_R_, restored its polymerase activity, whereas a substitution at position 3334, namely, BoDV-2 L_G_, did not (Fig. 2C). Accordingly, BoDV-2 L_RG_, which possesses both nucleotide substitutions simultaneously, exhibited the same level of polymerase activity as BoDV-2 L_R_ (Fig. 2C), indicating that a difference in the nucleotide at position 2762 is detrimental to the polymerase activity of BoDV-2 L.

To examine whether nucleotide differences at positions 2762 and 3334 of the L gene affect recovery of rBoDV-2 via reverse genetics, we constructed BoDV-2 cDNA plasmids expressing the full-length BoDV-2 antigenomes possessing each or both nucleotide substitutions. HEK293T cells were transfected with each BoDV-2 cDNA plasmid and five helper plasmids and cocultured with uninfected Vero cells at 3 days post transfection. After two weeks from coculture, rBoDV-2s that contain a nucleotide substitution at position 2762 of the L gene, rBoDV-2 L_R_ and L_RG_, were successfully rescued (Fig. 2D). Interestingly, although nucleotide substitution at position 3334 did not affect the polymerase activity of BoDV-2 L in the minireplicon assay (Fig. 2C), rBoDV-2 L_RG_, which possesses both substitutions at positions 2762 and 3334, seemed to be recovered more efficiently than rBoDV-2 L_R_, which possesses a substitution at position 2762 alone (Fig. 2D), suggesting that the nucleotide at position 3334 of the L gene also plays a role in BoDV-2 replication. To examine the growth ability of rBoDV-2 L_RG_ in comparison with that of the nonrecombinant BoDV-2 isolate No/98, Vero cells infected with rBoDV-2 L_RG_ or isolate No/98 were cocultured with uninfected Vero cells at a ratio of 1 to 19 (Fig. 2E), and viral propagation and the copy number of viral RNA were evaluated every 3 or 4 days by immunofluorescence assay (IFA) and RT-qPCR, respectively. While the nonrecombinant isolate No/98 spread to almost all Vero cells within 2 weeks of coculture, infection of rBoDV-2 L_RG_ exhibited limited spread within this period (Fig. 2F). Similarly, the copy number of viral RNA increased 20-fold in isolate No/98-infected cells, whereas it barely increased in rBoDV-2 L_RG_-infected cells (Fig. 2G). These results suggest that the growth ability of rBoDV-2 is not only determined by the L2762 and L3334 substitutions, even though both substitutions induce high polymerase activity in the minireplicon assay.

### Nonsynonymous SNPs in the L gene facilitate the growth of BoDV-2

Our RNA-seq analysis identified two nonsynonymous SNPs in the CTD of the L gene (Fig. 1A), indicating that the Vero cells persistently infected with BoDV-2 isolate No/98 contain a viral L quasispecies. Therefore, we constructed L expression plasmids harboring each or both nucleotide substitutions at positions 3743 and 4250 based on BoDV-2 L_RG_ and performed a minireplicon assay (Fig. 3A). While the substitution of uracil at position 3743 with cytosine, L_RGT_, increased polymerase activity by 1.3-fold, that of adenine at position 4250 with guanine, L_RGR_, decreased it by 0.7-fold. Substitution of both nucleotides, L_RGTR_, slightly decreased polymerase activity (Fig. 3B).

**Fig. 3.**
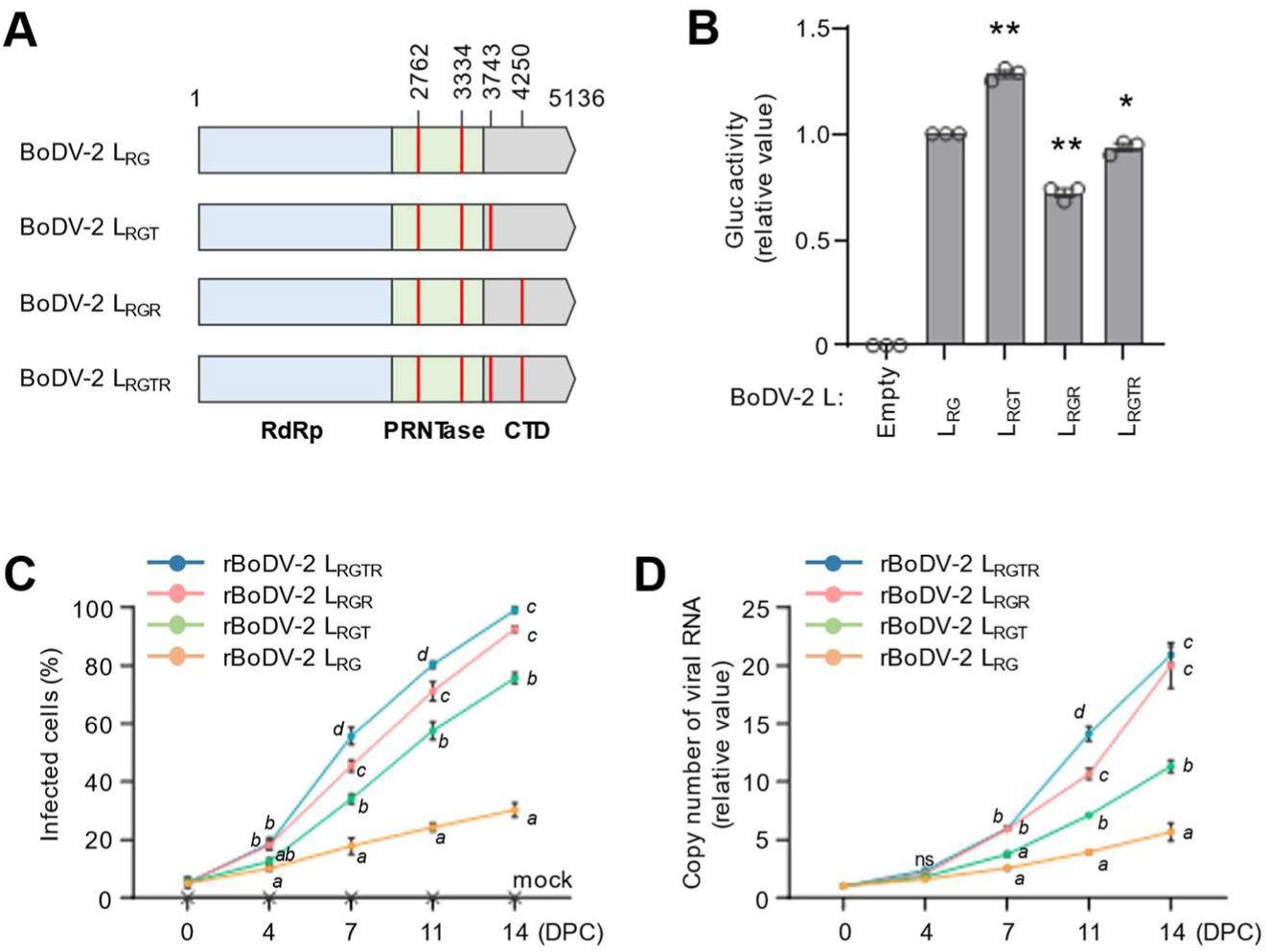
Nonsynonymous SNPs in the L gene facilitate the growth of BoDV-2. (A) Schematic representation of the BoDV-2 L gene variants inserted into the expression plasmid and the full-length BoDV-2 cDNA plasmid. The positions of the substituted nucleotides compared to the reference sequence are indicated by red bars. The putative RdRp domain, PRNTase domain, and CTD are colored blue, green, and gray, respectively. (B) BoDV-2 minireplicon assay using the indicated BoDV-2 L variants and BoDV-2 N and P as helper plasmids. Values for Gluc activity were normalized to that of the L_RG_ variant as a relative value of 1.0. Data are presented as the means ± SEM of three biologically independent experiments. One-way ANOVA and Dunnett’s multiple comparisons test were performed for statistical analysis. *, *p* < 0.05; **, *p* < 0.01. (C and D) Growth kinetics assay of rBoDV-2 variants. Vero cells infected with the indicated rBoDV-2 were cocultured with uninfected Vero cells at a ratio of 1 to 19 and passaged every 3 or 4 days. DPC, days post coculture. (C) The percentage of infected cells was determined by IFA using an antibody against BoDV N. (D) The copy number of viral RNA was measured via RT-qPCR using total RNA extracted from each cells. Data are presented as the means ± SEM of three biologically independent experiments. Two-way ANOVA and Tukey’s multiple comparisons test were performed for statistical analysis. Different characters (a, b, c) represent statistically significant differences at *p* < 0.05; ns, not significant.

To further investigate the impact of these SNPs on the growth ability of BoDV-2, we generated rBoDV-2 possessing each or both substitutions in the L gene and evaluated their growth kinetics. Although the substitution at position 4250 of the L gene had a detrimental effect on polymerase activity in the minireplicon assay (Fig. 3B), L proteins possessing either SNP substitution remarkably facilitated viral propagation and RNA synthesis of rBoDV-2 (Fig. 3C and D). These findings suggest that these SNPs influence viral replication through mechanisms other than enhancing polymerase activity in cells.

### A nonsynonymous SNP in the P gene expedites the growth of BoDV-2

We also identified another nonsynonymous SNP at position 456 of the P gene (Fig. 1A). The P gene encodes the viral polymerase cofactor, which directly interacts with the L protein and plays a crucial role in viral polymerase activity (24, 25). Therefore, we examined the effects of this SNP on viral polymerase activity and viral growth ability using both a minireplicon assay and rBoDV-2 variants. Substitution of guanine at position 456 of the P gene with adenine, P_I_, resulted in a slight increase in the polymerase activity of L_RG_ but not that of L_RGTR_, as observed in the minireplicon assay (Fig. 4A and B). Furthermore, the P gene with the introduced SNP significantly facilitated viral propagation (Fig. 4C and D) and RNA synthesis of rBoDV-2 (Fig. 4E and F). These observations indicate that the nonsynonymous SNP detected in the P gene also contributed to the increase in rBoDV-2 growth ability.

**Fig. 4.**
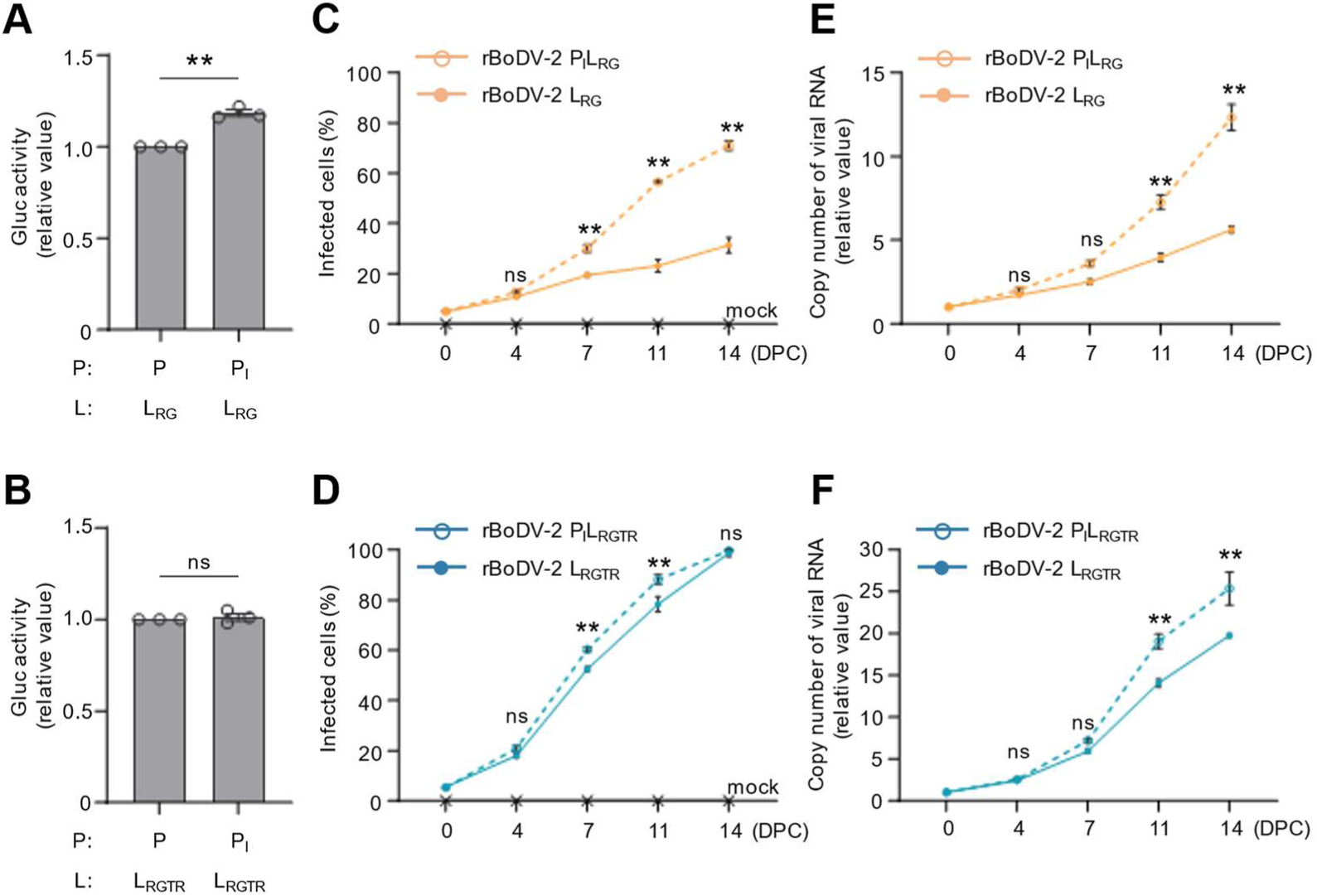
A nonsynonymous SNP in the P gene expedites the growth of BoDV-2. (A and B) BoDV-2 minireplicon assays using the indicated BoDV-2 P variants and BoDV-2 N and (A) L_RG_ or (B) L_RGTR_ as helper plasmids. Data are presented as the means ± SEM of three biologically independent experiments. An unpaired *t*-test was performed for statistical analysis. ns, not significant; **, *p* < 0.01. (C to F) Growth kinetics assay of rBoDV-2 variants. Vero cells infected with the indicated rBoDV-2 were cocultured with uninfected Vero cells at a ratio of 1 to 19 and passaged every 3 or 4 days. DPC, days post coculture. (C and D) The percentage of infected cells was determined by IFA using an antibody against BoDV N. (E and F) The copy number of viral RNA was measured via RT-qPCR using total RNA extracted from each cells. The data are presented as the means ± SEM of three biologically independent experiments. Two-way ANOVA and Tukey’s or Sidak’s multiple comparisons test were performed for statistical analysis. **, *p* < 0.01; ns, not significant.

### Lack of superinfection exclusion maintains the low-fitness polymorphism in cells persistently infected with BoDV-2

Since the isolation of BoDV-2 isolate No/98 from a pony brain in 1999, it has been maintained for a long time in persistently infected Vero cells. Notably, although BoDV-2-infected Vero cells have undergone repeated passages, L- and P-sequence variants with low viral replication ability appear to be the major quasispecies within persistently infected cells. The coexistence of viral quasispecies in infected cells has been shown to affect overall viral replication and pathogenicity and therefore plays a role in the adaptation and evolution of viral populations (26).

To determine the significance of the coexistence of BoDV-2 quasispecies in persistently infected cells, we performed infection experiments using cells infected with different sequences of rBoDV-2. First, we generated rBoDV-2 with different fluorescent marker genes, mCherry and GFP, inserted into the artificial intergenic region between the P and M genes (Fig. 5A) based on rBoDV-2 P_I_L_RGTR_ with high-growth ability. As shown in Fig. 5B, both viruses exhibited similar growth kinetics, confirming that the effects of these two fluorescence marker proteins on the growth ability of BoDV-2 are not different. We therefore examined the propagation of rBoDV-2 variants with different growth abilities, namely, low-growth rBoDV-2 L_RG_-mCherry (L_RG_-mCherry) and high-growth rBoDV-2 P_I_L_RGTR_-GFP (P_I_L_RGTR_-GFP), by a competition assay using Vero cells infected with each recombinant virus. When Vero cells infected with L_RG_-mCherry or P_I_L_RGTR_-GFP were cocultured with uninfected Vero cells at a ratio of 1:1:23, P_I_L_RGTR_-GFP rapidly spread throughout the culture, and the number of mCherry-expressing cells gradually decreased (Fig. 5C). Similarly, after 14 days of coculture, the genomic RNA of P_I_L_RGTR_-GFP accounted for almost all the BoDV-2 genomic RNA in the population (Fig. 5D). However, at 14 days after coculture, almost 1.0% of the cells coexpressed mCherry and GFP, and the proportion of cells expressing mCherry alone or coexpressing mCherry and GFP was approximately 3.3% of the total (Fig. 5C), indicating that high-growth P_I_L_RGTR_-GFP superinfected L_RG_-mCherry-infected cells and that low-growth rBoDV-2 was maintained without elimination from the cells superinfected with the P_I_L_RGTR_-GFP.

**Fig. 5.**
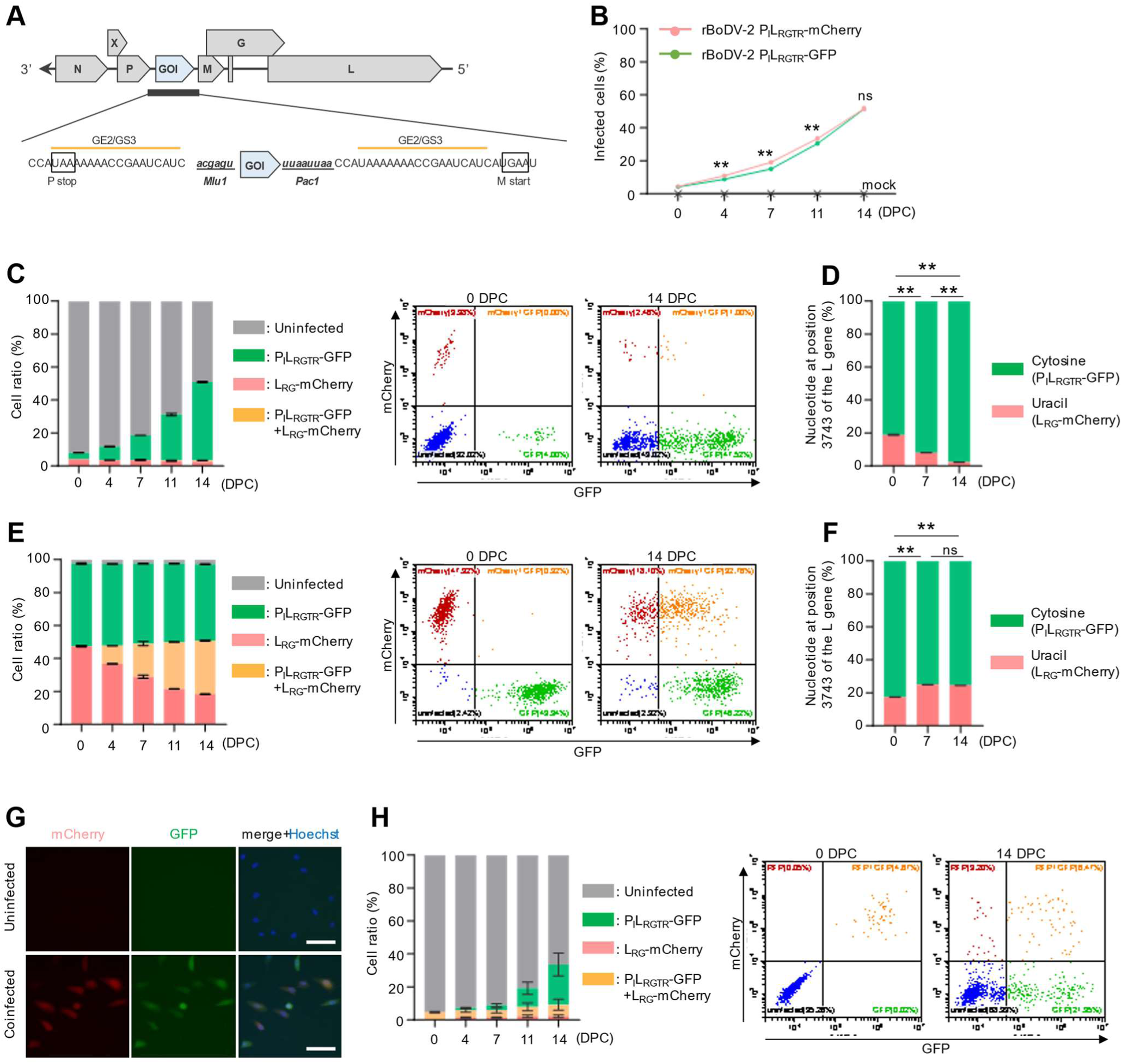
Lack of superinfection exclusion maintains the low-fitness polymorphism in cells persistently infected with BoDV-2. (A) Schematic representation of rBoDV-2 harboring an artificial expression cassette for the gene of interest (GOI) between the P and M genes. (B) Growth kinetics assay of rBoDV-2 harboring a fluorescent protein. Vero cells infected with rBoDV-2 P_I_L_RGTR_-mCherry or rBoDV-2 P_I_L_RGTR_-GFP were cocultured with uninfected Vero cells at a ratio of 1 to 19 and passaged every 3 or 4 days. The ratio of cells expressing fluorescent proteins was measured via flow cytometry. DPC, days post coculture. Data are presented as the means ± SEM of three biologically independent experiments. Two-way ANOVA and Sidak’s multiple comparisons test were performed for statistical analysis. **, *p* < 0.01; ns, not significant. (C and D) Competition assay between rBoDV-2 L_RG_-mCherry and rBoDV-2 P_I_L_RGTR_-GFP. The rBoDV-2 L_RG_-mCherry-infected, rBoDV-2 P_I_L_RGTR_-GFP-infected, and uninfected Vero cells were cocultured at a ratio of 1 to 1 to 23 and passaged every 3 or 4 days. DPC, days post coculture. (C) The ratio of cells expressing fluorescent proteins was measured via flow cytometry. Representative data at 0 and 14 DPC are shown. (D) A nucleotide at position 3743 of the L gene was analyzed via amplicon sequencing, and the ratios of uracil to cytosine, which represent rBoDV-2 L_RG_-mCherry and rBoDV-2 P_I_L_RGTR_-GFP, respectively, were compared. (E and F) Competition assay between rBoDV-2 L_RG_-mCherry and rBoDV-2 P_I_L_RGTR_-GFP. The rBoDV-2 L_RG_-mCherry-infected and rBoDV-2 P_I_L_RGTR_-GFP-infected Vero cells were cocultured at a ratio of 1 to 1 and passaged every 3 or 4 days. DPC, days post coculture. (E) The ratio of cells expressing fluorescent proteins was measured via flow cytometry. Representative data at 0 and 14 DPC are shown. (F) A nucleotide at position 3743 of the L gene was analyzed via amplicon sequencing, and the ratios of uracil to cytosine, which represent rBoDV-2 L_RG_-mCherry and rBoDV-2 P_I_L_RGTR_-GFP, respectively, were compared. (G) Isolation of Vero cells coinfected with rBoDV-2 L_RG_-mCherry and rBoDV-2 P_I_L_RGTR_-GFP. Bar, 500 μm. (H) Growth kinetics assay of rBoDV-2. Vero cells coinfected with rBoDV-2 L_RG_-mCherry and rBoDV-2 P_I_L_RGTR_-GFP were cocultured with uninfected Vero cells at a ratio of 1 to 19 and passaged every 3 or 4 days. DPC, days post coculture. The ratio of cells expressing fluorescent proteins was measured via flow cytometry and representative data at 0 and 14 DPC are shown. (C, E, and H) Data are presented as the means ± SEM of three biologically independent experiments. (D and F) Data are presented as the means ± SEM of three biologically independent experiments. Two-way ANOVA and Sidak’s multiple comparisons test were performed for statistical analysis. **, *p* < 0.01; ns, not significant.

To confirm the absence of superinfection exclusion between the rBoDV-2 variants, we next cocultured Vero cells infected with L_RG_-mCherry and P_I_L_RGTR_-GFP at a 1:1 ratio. As a result, the percentage of cells coexpressing mCherry and GFP gradually increased; after 14 days of coculture, the proportions of cells expressing GFP alone, mCherry alone, and coexpressing mCherry and GFP were approximately 46.3%, 18.3%, and 32.4%, respectively (Fig. 5E). This finding suggests that the high-growth P_I_L_RGTR_-GFP predominantly superinfected to the L_RG_-mCherry-infected cells. On the other hand, the proportion of viral genomic RNA of L_RG_-mCherry in the culture increased to some extent (Fig. 5F), indicating that the resident viruses were not eliminated even among the cells superinfected with the high-growth virus; rather, the replication of the low-growth viral genome was upregulated by support from the polymerase derived from high-growth virus. In addition, we cloned Vero cells coinfected with both L_RG_-mCherry and P_I_L_RGTR_-GFP (Fig. 5G) and cocultured them with uninfected Vero cells to investigate whether superinfection affects the propagation abilities of coinfected variants. As shown in Fig. 5H, the proportion of cells infected with only the P_I_L_RGTR_-GFP virus increased; the L_RG_-mCherry virus could also be transmitted to uninfected Vero cells, albeit only slightly. These observations suggest that BoDV-2 does not exclude superinfection, allowing low-fitness variants to persist within infected cells while maintaining their characteristics and thereby maintaining population diversity.

## Discussion

In this study, to better understand the virological characteristics of BoDV-2, we reassessed the whole-genome sequence of BoDV-2 using RNA extracted from persistently BoDV-2-infected Vero cells. Compared to the original sequence of BoDV-2 isolate No/98, the reassessed sequence contains two nucleotide substitutions at positions 2762 and 3334 of the L gene, both of which induce an amino acid change in the PRNTase domain (Fig. 1A and B). We found that substitution of uracil at position 2762 with guanine alone is sufficient to restore the polymerase activity of BoDV-2 L and to recover rBoDV-2 by reverse genetics (Fig. 2). This nucleotide substitution induces an amino acid change from leucine to arginine at position 921 of L. Interestingly, arginine is completely conserved at the same position in all 47 sequences of BoDV-1 L registered in the database, suggesting that this residue is critical for the polymerase activity of BoDV L. In contrast, the substitution of uracil at position 3334 with guanine, which induces an amino acid change from cysteine to glycine at position 1112 of L, did not affect the polymerase activity of BoDV-2 L in the minireplicon assay (Fig. 2C). However, interestingly, rBoDV-2, which possesses substitutions at positions 2762 and 3334, rBoDV-2 L_RG_, seemed to be rescued more efficiently than was rBoDV-2, which possesses a substitution at position 2762 alone, rBoDV-2 L_R_, by reverse genetics (Fig. 2D). This observation suggests that substitution at position 3334 may also play a critical role in the replication of BoDV-2 in a way other than enhancing polymerase activity. Position 3334 is located between motifs C (W) and D (HR) in the PRNTase domain, which is associated with the viral mRNA capping reaction (27), but it overlaps with neither these conserved motifs nor the currently identified signal sequences in BoDV-1 L (28, 29). Further investigations are needed to determine the effect of guanine at position 3334 of BoDV-2 L on viral propagation.

Our study showed that while rBoDV-2 could be rescued by substituting nucleotides at positions 2762 and 3334 of the L gene, the growth ability of rBoDV-2 L_RG_ was severely attenuated compared to that of the nonrecombinant BoDV-2 isolate No/98 (Fig. 2F and G). Interestingly, the substitution of nonsynonymous SNPs at positions 3743 and 4250 of the L gene significantly facilitated the growth ability of rBoDV-2 (Fig. 3C and D). Nucleotide substitution of uracil at position 3743 with cytosine induces an amino acid change from isoleucine to threonine at amino acid position 1248 of BoDV-2 L. Either isoleucine or threonine is conserved at the same position in BoDV-1 Ls in the database at a similar ratio, though it remains unclear whether substitution of isoleucine with threonine also affects the viral growth ability of BoDV-1. In addition, nucleotide substitution of alanine at position 4250 with guanine induces an amino acid change from lysine to arginine at amino acid position 1417 of BoDV-2 L. All the BoDV-1 Ls in the database except for that of strain CRNP5, which was generated by several passages of isolate He/80 in the brains of Lewis rats and SJL mice, show conservation of a lysine residue at the same amino acid position corresponding to 1417 of BoDV-2 L (30), but it is unclear whether rBoDV-1 possessing arginine residue at this position shows efficient propagation compared to the parental BoDV-1 isolate He/80 (31). Investigating how these SNPs affect viral replication may help to elucidate the previously unknown role of the CTD of BoDV L.

In this study, we also identified one nucleotide substitution within an SNP at position 456 of the P gene. This substitution leads to an amino acid change from methionine to isoleucine at position 152, which facilitates the growth ability of rBoDV-2 (Fig. 4). BoDV-1 P interacts with P and L through binding domains spanning amino acid positions 135 to 172 and 135 to 183, respectively (25, 32). Additionally, previous research has indicated the presence of a nuclear export signal between amino acid positions 145 and 165 within BoDV-1 P (33). Although the role of the amino acid residue in BoDV-1 P, corresponding to position 152 in BoDV-2 P, has yet to be explored, investigating its potential binding interactions with other viral proteins and its influence on the nuclear translocation capability of P would be of considerable interest.

Detection of 3 major nonsynonymous SNPs suggests the presence of up to eight distinct BoDV-2 genomes within persistently infected Vero cells. BoDV-2 was initially isolated from the brain of an infected pony using primary young rabbit brain cells and subsequently propagated in Vero cells (5), after which the infected Vero cells were maintained for two decades. It is unclear when and how BoDV-2 acquired the P and L gene SNPs that we found in this study. However, given the comparable proportion of genome sequences containing SNPs, even those associated with low-fitness effects on propagation in culture, it may be possible that these SNPs were acquired after isolation *in vitro* to maintain persistent infection within cultured cells. On the other hand, our analysis revealed that BoDV-2 does not exhibit superinfection exclusion, even toward low-fitness mutant viruses (Fig. 5). From this observation, it is conceivable that viruses with different sequences may have coexisted before isolation. To better understand the characteristics and evolution of BoDV-2 infection, it is necessary to examine in detail the competition for growth between viruses with genomic variants not only *in vitro* but also *in vivo*.

In this study, we demonstrated that rBoDV-2 can infect cells infected with another rBoDV-2 with different replication abilities (Fig. 5C and E), showing that BoDV-2 does not exhibit superinfection exclusion. Superinfection exclusion is a common strategy used by many viruses to prevent sequential infection by closely related viruses (34, 35). For instance, substantial infection of influenza A virus (IAV) in cells that are already infected with IAV is inhibited by the active viral replication complex (36). It has been reported that IAV superinfection exclusion constrains interactions between viral populations locally within a host to regulate viral fitness and evolution (37). While superinfection exclusion provides evolutionary advantages for controlling viral fitness and population diversity, it also has the effect of promoting founder effects and reducing diversity and is thought to control the balance between genetic diversification and genome integrity in infected hosts (26). A simulation study demonstrated that although a mutant with superinfection exclusion can overtake a population that cannot exclude superinfection, superinfection enables a population to adapt to environmental changes, indicating that superinfection exclusion negatively affects the adaptation of a viral population in the long term (38). These observations suggest that maintenance of genetic diversity through the absence of superinfection exclusion during BoDV-2 infection plays a pivotal role in the establishment of persistent infection. This might be achieved by not excluding low-fitness variants that may regulate replication dynamics and cytopathic effects as a viral population in infected cells.

In previous studies, BoDV-1 superinfection in cells persistently infected with BoDV-1 was demonstrated to be excluded both *in vivo* and *in vitro* (39, 40). The authors cocultured UTA6 human osteosarcoma cells persistently infected with the BoDV-1 isolate He/80 with either uninfected or BoDV-1 isolate H215-infected Vero cells and evaluated infection of isolate He/80 to Vero cells by IFA using a monoclonal antibody that reacts with the N of He/80 but not with the N of H215. Although infection of the isolate He/80 spread efficiently from the UTA6 cells to uninfected Vero cells, the infection did not spread to the H215-infected Vero cells after 3 weeks of coculture (40). The authors further demonstrated that expression of either N, accessory protein (X), or P is sufficient to inhibit superinfection with BoDV-1, concluding that the imbalance in intracellular levels of vRNP components is attributable to superinfection exclusion of BoDV-1 (40). However, whereas the growth abilities of both the BoDV-1 isolates H215 and He/80 were high enough for the infection to spread efficiently in the previous study, the growth ability of the two different rBoDV-2s was significantly different in our experiments (Fig. 5C). In addition, the methods used to detect superinfection differed between these studies. Thus, further studies are necessary to investigate whether the absence of superinfection exclusion is a specific property for BoDV-2 but not for BoDV-1.

In conclusion, we successfully established a reverse genetics system for BoDV-2 and revealed lack of superinfection exclusion enables BoDV-2 to maintain the genetic diversity in persistently infected cells. Our study provides valuable insights into the biological properties of BoDV-2 and contributes to development of effective vaccines and antiviral drugs for pathogenic mammalian orthobornaviruses.

## Materials and Methods

### Cell culture

Human embryonic kidney (HEK) 293T cells were cultured in Dulbecco’s modified Eagle’s medium (DMEM) (Nacalai Tesque, Japan; #08456-36) supplemented with 10% fetal bovine serum (FBS) (Thermo Fisher Scientific, USA; #10270-106) and 1% penicillin-streptomycin-amphotericin B solution (Fujifilm, Japan; #161-23181). African green monkey kidney Vero cells were cultured in DMEM supplemented with 5% FBS and 1% penicillin-streptomycin-amphotericin B solution.

### Virus

Vero cells persistently infected with BoDV-2 isolate No/98 were kindly provided by Prof. Martin Schwemmle (University of Freiburg, Germany) and propagated by Dennis Rubbenstroth’s group at the Friedrich-Loeffler-Institut (FLI), Greifswald, Germany. Isolate No/98 had originally been isolated using primary young rabbit brain cells, which were subsequently cocultured with Vero cells. After consecutive passages, only Vero cells remained in the culture (5). The exact propagation history of the isolate No/98-infected culture is unknown.

### RNA-seq and data processing

Total RNA was extracted from persistently BoDV-2-infected Vero cells using Agencourt® RNAdvance Tissue kit (Beckman Coulter) and the KingFisher Flex system (Thermo Fisher Scientific) according to manufacturers’ instructions. Subsequent cDNA synthesis and library preparation were performed as described in Wylezich et al. (48) with slight modifications. Briefly, cDNA was synthesized using the SuperScript IV First-Strand cDNA Synthesis System (Thermo Fisher Scientific) and subsequently, the complete reaction was used for 2nd strand synthesis with the NEBNext Ultra II Non-Directional RNA Second Strand Synthesis Module (New England Biolabs), followed by DNA fragmentation with a Covaris M220 and library preparation with a GeneRead DNA Library L Core kit (QIAGEN) and ION Xpress Barcode adapters (Thermo Fisher Scientific). After size selection as described in (48), library size was measured using a 2100 BioAnalyzer system (Agilent Technologies) with Agilent High Sensitivity DNA Chip and reagents, and the library was quantified using a QIAseq Library Quant Assay Kit (Qiagen). The library was prepared for sequencing and sequenced together with Ion Torrent Calibration Standard (Thermo Fisher) using an Ion Torrent S5 XL instrument with Ion 530 chip and reagents (Thermo Fischer Scientific) in 400 bp mode. Sequencing adapters were trimmed using the Newbler assembler of the 454 Genome Sequencer Software Suite v. 3.0 (Roche) and the FastQC was used to quality control the sequence dataset. Initial reference-based mapping (454 Genome Sequencer Software Suite v. 3.0, Roche) against RefSeq BoDV-2 strain No/98 complete genome (NC_030692) was performed to generate a sample-specific consensus sequence. This consensus sequence was used as a reference for the second round of reference-based mapping analysis. Pairwise sequence alignment of the resulting BoDV-2 genome sequence and BoDV-2 AJ311524.1 was performed MUSCLE (49). This alignment was visually inspected with Geneious Prime® 2019.2.3 Software (Biomatters). SNPs within the BoDV-2 genome were detected using the Find Variations/SNPs program implemented in the Geneious Prime® 2019.2.3 software. For this analysis, we used the BAM file generated from the 2nd round of mapping analyses as data input and the following parameters, minimum coverage: 50, minimum variant frequency: 25%, maximum variant *p*-value 10 (raise to) -7, minimum strand bias *p*-value of 10 (raise to) -5 when exceeding 65% bias.

### RACE analysis

Total RNA extracted from BoDV-2-infected Vero cells was ligated with either 5’ adaptor oligoRNA (5’-CGACUGGAGCACGAGGACACUGACAUGGACUGAAGGAGUAGAAA (Thermo Fisher Scientific, USA; #L150201)) or 3’ adaptor oligoRNA (5’-GAAGAGAAGGUGGAAAUGGCGUUUUGG; the 5’ and 3’ ends were modified with a monophosphate and an amino-modifier, respectively) using T4 RNA Ligase 1 (New England Biolabs, USA; #M0204) according to the manufacturer’s instructions. The adaptor-ligated RNA was reverse transcribed using SuperScript Ⅳ Reverse Transcriptase (Thermo Fisher Scientific, USA; #18090010) and amplified by PCR using Q5 Hot Start High-Fidelity 2x Master Mix (New England Biolabs, USA; #M0494) according to the manufacturer’s instructions. The viral terminal sequences were analyzed via Sanger sequencing of the purified PCR amplicons. The primers used in the RACE analysis are listed in the Table. 1.

**Table. 1.**
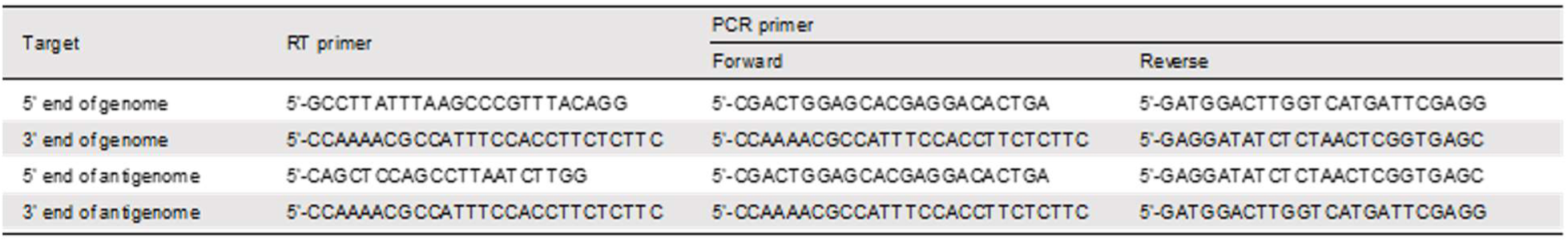

### Plasmid construction

The plasmid used to express BoDV-2 N, P, and L were described previously (23). Nucleotide substitutions in the P or L gene were introduced by PCR mutagenesis. A FLAG-tag sequence was inserted with a GS-linker sequence into the 5’ terminus of the L gene by PCR mutagenesis. A plasmid expressing each BoDV-2 M or G was constructed by inserting each gene into the EcoRⅠ and XhoⅠ restriction enzyme recognition sites of the eukaryotic expression plasmid pCAGGS (41). For both plasmids, synonymous substitutions were introduced by PCR mutagenesis into the splicing donor and acceptor sites to increase protein expression levels (20, 42). An RNA polymerase Ⅱ (Pol Ⅱ)-driven BoDV-2 minigenome plasmid was constructed similarly to the BoDV-1 minireplicon plasmid described previously (23). In brief, a *Gaussia* luciferase (Gluc) reporter gene flanked by the 3’ and 5’ untranslated regions of BoDV-2 in the negative-sense orientation was inserted into the XhoⅠ and NotⅠ restriction enzyme recognition sites of pCAG-HRSV3 (43), with a hammerhead ribozyme (HamRz) sequence at the 5’ end and a hepatitis delta virus ribozyme (HdvRz) sequence at the 3’ end. A BoDV-2 cDNA plasmid that expresses the full-length antigenomic RNA of BoDV-2 strain No/98 was constructed similarly to the corresponding BoDV-1 cDNA plasmid described previously (44). In brief, an artificially synthesized full-length cDNA identical to the published reference sequence of BoDV-2 isolate No/98 was inserted into the XhoⅠ and NotⅠ restriction enzyme recognition sites of pCAG-HRSV3 (43) with an HamRz sequence at the 5’ end and an HdvRz sequence at the 3’ end. A BoDV-2 cDNA plasmid that harbors an additional expression cassette between the P and M genes was constructed by inserting a gene of interest flanked by the MluⅠ and PacⅠ restriction enzyme recognition sequences with the repeated 2nd gene end (GE2) and the 3rd gene start (GS3) signal sequences by PCR mutagenesis (Fig. 5A) (44).

### Western blotting

Cultured cells were lysed with 1x sample buffer (50 mM Tris-HCl, pH 6.8; 2% sodium dodecyl sulfate (SDS); 5% w/v sucrose; and 5% 2-mercaptoethanol) followed by boiling at 95 °C for 10 minutes. Cell lysates were subjected to SDS-PAGE using an ePAGEL (ATTO Corporation, Japan; #2331720), and proteins were subsequently transferred to Trans-Blot® Turbo™ Mini nitrocellulose membranes (Bio-Rad, USA; #1704156). The membranes were blocked with blocking buffer (TBS containing 0.1% Tween-20 with 5% w/v skim milk) and then incubated with the primary antibody (anti-FLAG M2 (Sigma-Aldrich, USA; #F1804) or anti-alpha-tubulin (Sigma-Aldrich, USA; #T5168)), followed by incubation with the secondary antibody (horseradish peroxidase (HRP)-conjugated anti-mouse IgG (Jackson ImmunoResearch, USA; #715-035-150). The protein bands were detected with Clarity Western ECL substrate reagents (Bio-Rad, USA; #170-5060), and chemiluminescence signals were visualized by Fusion Solo S (Vilber, France).

### BoDV-2 minireplicon assay

HEK293T cells seeded into 48-well plates were transfected with 50 ng of BoDV-2 minigenome plasmid, 50 ng of pCAGGS-BoDV-2 N, 5 ng of pCAGGS-BoDV-2 P, 50 ng of pCAGGS-BoDV-2 L, and 5 ng of control plasmid expressing *Cypridina* luciferase using TransIT293 (Mirus, USA; #MIR2700) according to the manufacturer’s instructions. At 48 hours post transfection, *Gaussia* luciferase and *Cypridina* luciferase expression levels were measured with a Biolux *Gaussia* luciferase assay kit (New England Biolabs, USA; #E3300) and Biolux *Cypridina* luciferase assay kit (New England Biolabs, USA; #E3309), respectively, according to the manufacturer’s instructions. Luminescence signals were detected by a GloMax Discover Microplate Reader (Promega, USA), and polymerase activity was quantified by normalizing *Gaussia* luciferase activity to *Cypridina* luciferase activity.

### Reverse genetics of BoDV-2

Reverse genetics was performed according to the protocol summarized in our previous review article (21). HEK293T cells seeded into 6-well plates were transfected with 2.0 µg of BoDV-2 cDNA plasmid, 0.5 µg of pCAGGS-BoDV-2 N, 0.025 µg of pCAGGS-BoDV-2 P, 0.25 µg of pCAGGS-BoDV-1 L, 0.04 µg of pCAGGS-BoDV-2 M, and 0.01 µg of pCAGGS-BoDV-2 G using TransIT293 according to the manufacturer’s instructions. At 3 days post transfection, the transfected cells were cocultured with uninfected Vero cells, which were passaged every 3 or 4 days until the infection spread to almost all the Vero cells. The transfected HEK293T cells were removed by adding 1.0 μg/mL puromycin (InvivoGen, USA; #ant-pr) at the optimal time point.

### Immunofluorescence assay

Cultured cells were fixed with 4% paraformaldehyde (Nacalai Tesque, Japan; #09154-85) for 10 minutes and permeabilized with PBS containing 0.5% Triton X-100 (Wako, Japan; #169-21105) and 5% bovine serum albumin (Sigma-Aldrich, USA; #A3059) for 10 minutes. The cells were incubated with the primary antibody (polyclonal rabbit serum anti-BoDV N HB01), followed by incubation with the secondary antibody (Alexa Fluor® 555-conjugated anti-rabbit IgG (Thermo Fisher Scientific, USA; #A21429)) and 300 nM 4′,6′-diamidino-2-phenylindole (DAPI) (Merck, Germany; #28718-90-3). Fluorescence signals were observed by an ECLIPSE TE2000-U fluorescence microscope (Nikon, Japan).

### RT-qPCR

Total RNA was extracted from cultured cells using NucleoSpin RNA Plus (Macherey-Nagle, Germany; #740984) according to the manufacturer’s instructions. For detection of BoDV-2 RNA, RT-qPCR was performed using qScript XLT one-step RT-qPCR ToughMix Kit (Quanta Biosciences, USA; #95132) with forward primer (5’-TAGTYAGGAGGCTCAATGGCA), reverse primer (5’-GTCCYTCAGGAGCTGGTC), and probe (5’-FAM-AAGAAGATCCCCAGACACTACGACG-BHQ1). Each reaction contained 2.75 μl of RNase-free water, 6.25 μl of 2x qScript XLT One-Step RT-qPCR ToughMix, 1.0 μl of primer-probe mix (7.5 pmol/μl primer and 2.5 pmol/μl probe) and 2.5 μl of RNA template (approximately 500 ng of total RNA) or RNase-free water for the no template control in a total volume of 12.5 μl. The thermal program consisted of 1 cycle of 50 °C for 10 min and 95 °C for 1 min, followed by 40 cycles of 95 °C for 10 sec, 57 °C for 30 sec, and 68 °C for 30 sec (8). For detection of GAPDH mRNA, approximately 500 ng of total RNA was reverse transcribed using the Verso cDNA synthesis kit (Thermo Fisher Scientific, USA; #AB1453) with anchored oligo dT primer according to the manufacturer’s instructions. qPCR was performed using Luna Universal qPCR Master Mix (New England Biolabs, USA; #M3003) with forward (5’-ATCTTCTTTTGCGTCGCCAG) and reverse (5’-ACGACCAAATCCGTTGACTCC) primers. Each reaction contained 8.0 μl of RNase-free water, 10.0 μl of Luna Universal qPCR Master Mix, 1.0 μl of primer mixture (10.0 pmol/μl primer) and 1.0 μl of synthesized cDNA or RNase-free water for the no template control in a total volume of 20.0 μl. The thermal program consisted of 1 cycle of 95 °C for 1 min and 40 cycles of 95 °C for 10 sec and 60 °C for 30 sec, followed by a melting reaction. All reactions were performed with a CFX Connect real-time system (Bio-Rad, USA), and relative expression levels of BoDV-2 RNA were calculated via the relative quantification method with GAPDH mRNA serving as a reference.

### Flow cytometry

Cultured cells were fixed with 4% paraformaldehyde for 20 minutes and suspended in PBS containing 2.0% FBS. Proportions of cells expressing either or both mCherry and GFP were analyzed by a CytoFLEX S flow cytometer (Beckman Coulter, USA). The criterion for assessing the extent of positive detection of each mCherry or GFP signal was set using mock-infected cells as a negative control.

### Amplicon sequencing

Total RNA was extracted from cultured cells using NucleoSpin RNA Plus according to the manufacturer’s instructions. BoDV-2 genomic RNA was reverse transcribed using SuperScript Ⅳ Reverse Transcriptase with a genome-specific primer (5’-AGGGACACTCTCGTTCTCCA) according to the manufacturer’s instructions. To prepare the sequencing library, the 1st PCR product ranging from position 3556 to 3954 of the L gene was amplified from 1.0 μl of cDNA using Q5 Hot Start High-Fidelity 2x Master Mix with forward (5’-acactctttccctacacgacgctcttccgatctGGATGCCTCTATGCTGACTCTG) and reverse (5’-gtgactggagttcagacgtgtgctcttccgatctACCAACCAGGAATCGCCAAA) primers. Then, amplicon sequencing was performed by the Bioengineering Lab. Co., Ltd. Briefly, the 2nd PCR product was amplified from 10 ng purified 1st PCR product using the TaKaRa Ex Taq HS (TaKaRa Bio Inc., Japan; #RR006A) with a forward (5’-AATGATACGGCGACCACCGAGATCTACAC-*Index2*-ACACTCTTTCCCTACACGACGC) and reverse (5’-CAAGCAGAAGACGGCATACGAGAT-*Index1*-GTGACTGGAGTTCAGACGTGTG) primers. Sequencing of the purified 2nd PCR products was performed using the MiSeq system and MiSeq Reagent Kit v3 (Illumina, USA) with a 2x300 bp read length protocol. Adaptor and primer sequences were removed from the raw sequence reads using the FASTX Toolkit (ver. 0.0.14) (45). Low-quality and short reads were also removed using Sickle (ver. 1.33) with criteria of a quality score of <Q20 and a read length of ≤40 bp (46). Filtered reads were merged using FLASH (ver. 1.2.11) with default settings (47). The number of BoDV-2 genotypes was counted by analyzing the nucleotide at position 3743 in each read.

### Statistical analysis

All statistical analyses were performed with GraphPad Prism 10 software. The tests used for each experiment are described in figure legends.

## Acknowledgments

We are grateful to Prof. Martin Schwemmle (University of Freiburg, Germany) for providing BoDV-2-infected Vero cells and to Andrea Aebischer, Doreen Schulz, Patrick Zitzow, and Kathrin Steffen (Friedrich-Loeffler-Institut, Germany) for their technical support. We also thank Prof. Masayuki Horie (Osaka Metropolitan University, Japan) for supporting bilateral research.

This study was supported in part by JSPS KAKENHI grants JP19J23468 (TK), JP22K20507 (TK), JP23K14083 (TK), JP19K22530 (KT), JP20H05682 (KT), and JP21K19909 (KT); JSPS Overseas Challenge Program for Young Researchers grant 202080194 (TK); JSPS Core-to-Core Program JPJSCCA20190008 (KT); Kaketsuken Research Grant (KT); the Joint Usage/Research Center Program on Institute for Life and Medical Sciences, Kyoto University (KT); Institute for Life and Medical Sciences Office of Directors’ Research Grants Program, Kyoto University (TK); and the German Federal Ministry of Education and Research, Zoonotic Bornavirus Consortium (ZooBoCo), grants 01KI1722 and 01KI2005 (DR, MB, and DH).

## Notes

### Competing Interest Statement

The authors have declared no competing interest.

